# Learning from heterogeneous data sources: an application in spatial proteomics

**DOI:** 10.1101/022152

**Authors:** Lisa M. Breckels, Sean Holden, David Wojnar, Claire M. Mulvey, Andy Christoforou, Arnoud Groen, Matthew W.B. Trotter, Oliver Kohlbacher, Kathryn S. Lilley, Laurent Gatto

## Abstract

Sub-cellular localisation of proteins is an essential post-translational regulatory mechanism that can be assayed using high-throughput mass spectrometry (MS). These MS-based spatial proteomics experiments enable us to pinpoint the sub-cellular distribution of thousands of proteins in a specific system under controlled conditions. Recent advances in high-throughput MS methods have yielded a plethora of experimental spatial proteomics data for the cell biology community. Yet, there are many third-party data sources, such as immunofluorescence microscopy or protein annotations and sequences, which represent a rich and vast source of complementary information. We present a unique transfer learning classification framework that utilises a nearest-neighbour or support vector machine system, to integrate heterogeneous data sources to considerably improve on the quantity and quality of sub-cellular protein assignment. We demonstrate the utility of our algorithms through evaluation of five experimental datasets, from four different species in conjunction with four different auxiliary data sources to classify proteins to tens of sub-cellular compartments with high generalisation accuracy. We further apply the method to an experiment on pluripotent mouse embryonic stem cells to classify a set of previously unknown proteins, and validate our findings against a recent high resolution map of the mouse stem cell proteome. The methodology is distributed as part of the open-source Bioconductor `pRoloc` suite for spatial proteomics data analysis.

**Abbreviations:**

LOPIT: Localisation of Organelle Proteins by Isotope Tagging
PCP: Protein Correlation Profiling
ML: Machine learning
TL: Transfer learning
SVM: Support vector machine
PCA: Principal component analysis
GO: Gene Ontology
CC: Cellular compartment
iTRAQ: Isobaric tags for relative and absolute quantitation
TMT: Tandem mass tags
MS: Mass spectrometry

## Introduction

Cell biology is currently undergoing a data-driven paradigm shift [1]. Molecular biology tools, imaging, biochemical analyses and omics technologies, enable cell biologists to track the complexity of many fundamental processes such as signal transduction, gene regulation, protein interactions and sub-cellular localisation [2]. This remarkable success, has resulted in dramatic growth in data over the last decade, both in terms of size and heterogeneity. Coupled with this influx of experimental data, databases such as UniProt [3] and the Gene Ontology [4] have become more information rich, providing valuable resources for the community. The time is ripe to take advantage of complementary data sources in a systematic way to support hypothesis- and data-driven research. However, one of the biggest challenges in computational biology is how to meaningfully integrate heterogeneous data; transfer learning, a paradigm in machine learning, is ideally suited to this task.

Transfer learning has yet to be fully exploited in computational biology. To date, various data mining and machine learning (ML) tools, in particular classification algorithms have been widely applied in many areas of biology [5]. A classifier is trained to learn a mapping between a set of observed instances and associated external attributes (class labels) which is subsequently used to predict the attributes on data with unknown class labels (unlabelled data). In transfer learning, there is a primary task to solve, and associated primary data which is typically expensive, of high quality and targeted to address a specific question about a specific biological system/condition of interest. While standard supervised learning algorithms seek to learn a classifier on this data alone, the general idea in transfer learning is to complement the primary data by drawing upon an auxiliary data source, from which one can extract complementary information to help solve the primary task. The secondary data typically contains information that is related to the primary learning objective, but was not primarily collected to tackle the specific primary research question at hand. These data can be heterogeneous to the primary data and are often, but not necessarily, cheaper to obtain and more plentiful but with lower signal-to-noise ratio.

There are several challenges associated with the integration of information from auxiliary sources. Firstly, if the primary and auxiliary sources are combined via straightforward concatenation the signal in the primary can be lost through dilution with the auxiliary due to the latter being more plentiful and often having lower signal-to-noise ratio (see S5 File, Figure 7 for an illustration). Feature selection can be used to extract the attributes with the most distinct signals, however the challenge still remains in how to combine this data in a meaningful way. Secondly, combining data that exist in different data spaces is often not straightforward and different data types can be sensitive to the classifier employed, in terms of classifier accuracy.

In one of the first applications of transfer learning Wu and Dietterich [6] used a *k*-nearest neighbours (*k*-NN) and support vector machine (SVM) framework for plant image classification. Their primary data consisted of high-resolution images of isolated plant leaves and the primary task was to determine the tree species given an isolated leaf. An auxiliary data source was available in the form of dried leaf samples from a Herbarium. Using a kernel derived from the shapes of the leaves and applying the transfer learning (TL) framework [6], they showed that when primary data is small, training with auxiliary data improves classification accuracy considerably. There were several limitations in their methods: firstly, the data in the *k*-NN TL classifier were only weighted by data source and not on a class-by-class basis, and, secondly in the SVM framework both data sources were expected to have the same dimensions and lie in the same space. We present an adaption and significant improvement of this framework and extend the usability of the method by (i) incorporating a multi-class weighting schema in the *k*-NN TL classifier, and (ii) by allowing the integration of primary and auxiliary data with different dimensions in the SVM schema to allow the integration of heterogeneous data types. We apply this framework to the task of protein sub-cellular localisation prediction from high resolution mass spectrometry (MS)-based data.

Spatial proteomics, the systematic large-scale analysis of a cell’s proteins and their assignment to distinct sub-cellular compartments, is vital for deciphering a protein’s function(s) and possible interaction partners. Knowledge of where a protein spatially resides within the cell is important, as it not only provides the physiological context for their function but also plays an important role in furthering our understanding of a protein’s complex molecular interactions e.g. signalling and transport mechanisms, by matching certain molecular functions to specific organelles.

There are a number of sources of information which can be utilised to assign a protein to a sub-cellular niche. These range from high quality data produced from experimental high-throughput quantitative MS-based methods (e.g. LOPIT [7] and PCP [8]) and imaging data (e.g. [9]), to freely available data from repositories and amino acid sequences. The former, in a nutshell, involves cell lysis followed by separation and fractionation of the subcellular structures as a function of their density, and then selecting a set of distinct fractions to quantify by mass-spectrometry. These quantitative protein profiles are representative of organelle distribution and hence are indicative of their subcellular localisation [10]. Based on the distribution of a set of known genuine organelle marker proteins, pattern recognition and ML methods can be used to match and associate the distributions of unknown residents to that of one of the markers. There is thus a reliance on reliable organelle markers and statistical learning methods for robust proteome-wide localisation prediction [11]. These approaches have been utilised to gain information about the sub-cellular location of proteins in several biological examples, such as *Arabidopsis* [7, 12, 13, 14, 15, 16], *Drosophila* [17], yeast [18], human cell lines [19, 20], mouse [8, 21] and chicken [22], using a number of algorithms, such as, SVMs [23], *k*-NN [15], random forest [24], naive Bayes [14], neural networks [25], and partial-least squares discriminant analysis [7], [17], [22].

Although application of computational tools to spatial proteomics is a recent development, the determination of protein localisation using *in silico* data is well-established (reviewed in [26, 27, 28]). Many methods have been developed to predict protein localisation from amino acid sequence features e.g. amino-acid composition information (e.g. [29, 30, 31, 32, 33, 34, 35]), localisation signals and motifs relevant to protein sorting (e.g. [36, 37, 38, 39, 40, 41, 42, 43]). Annotation-based prediction methods have also been widely used that use information about functional domains (e.g. [44, 45]), protein-protein interactions (e.g. [46, 47, 48]) and Gene Ontology (GO) [4] terms (e.g. [49, 50, 51, 52]). However, not all proteins in GO are reliably annotated; for example, according to the 2015 03 release of UniProtKB [3] the human, mouse, *Drosophila melanogaster* and *Arabidopsis thaliana* proteomes have less than 14%, 14%, 6% and 13% experimentally-verified Gene Ontology cellular compartment (GO CC) sub-cellular annotations, in each proteome respectively.

Despite improvements in generalisation accuracy of sequence- and annotation-based classifiers, a fundamental problem concerns the biological relevance and ultimate utility to cell biology of such predictors. Protein sequences and their annotation do not change according to cellular condition or cell type, whereas protein localisation can change in response to cellular perturbation. Furthermore, this type of data does not adequately describe the range of mechanisms via which a particular protein may reside in a particular organelle. Not all protein sequences contain motifs or exhibit compositional properties indicative of organelle residency. Despite the inherent limitations of using *in silico* data to predict dynamic cell- and condition-specific protein properties, transfer learning [6, 53, 50, 51, 52] may allow the transfer of complementary information available from these data to classify proteins in experimental proteomics datasets.

Here, we present a new transfer learning framework for the integration of heterogeneous data sources, and apply it to the task of sub-cellular localisation prediction from experimental and condition-specific MS-based quantitative proteomics data. Using the *k*-NN and SVM algorithms in a transfer learning framework we find that when given data from a high quality MS experiment, integrating data from a second less information rich but more plentiful auxiliary data source directly in to classifier training and classifier creation results in the assignment of proteins to organelles with high generalisation accuracy. Five experimental MS LOPIT datasets, from four different species, were employed in testing the classifiers. We show the flexibility of the pipeline through testing four auxiliary data sources; (1) Gene Ontology terms, (2) immunocytochemistry data [9], (3) sequence and annotation features, and (4) protein-protein interaction data [54]. The results obtained demonstrate that this transfer learning method outperforms a single classifier trained on each single data source alone and on a class-by-class basis, highlighting that the primary data is not diluted by the auxiliary data. This methodology forms part of the open-source open-development Bioconductor [55] `pRoloc` [56] suite of computational methods available for organelle proteomics data analysis.

## Results

Here, we have adapted a classic application of inductive transfer learning (TL) [6] using experimental quantitative proteomics data as the primary source and Gene Ontology Cellular Compartment (GO CC) terms as the auxiliary source. Using this TL approach, we have exploited auxiliary data to improve upon the protein localisation prediction from quantitative MS-based spatial proteomics experiments using (1) a class-weighted *k*-NN classifier, and (2) an SVM classifier in a TL framework. We also show the flexibility of the framework by using data from the Human Protein Atlas [57] and input sequence and annotation features from the YLoc [58, 59] web server, and protein-protein interaction data from the STRING database [54] as auxiliary data sources.

To assess classifier performance we employed the classic machine learning schema of partitioning our labelled data into training and testing sets, and used the testing sets to assess the strength of our classifiers. Parameter optimisation was conducted on the labelled training data using 100 rounds of stratified 80/20 partitioning, in conjunction with 5-fold cross-validation in order to estimate the free parameters via a grid search, as implemented in the `pRoloc` package [56] (and described in the methods below). Here, for the *k*-NN TL algorithm these parameters are the weights assigned to each class for each data source, and for the SVM TL algorithm these are *C*, *γP* and *γA* for the two kernels, as described in the materials and methods. The testing set is then used to assess the generalisation accuracy of the classifier. By applying the best parameters found in the training phase on test data, observed and expected classification results can be compared, and then used to assess how well a given model works by getting an estimate of the classifier’s ability to achieve a good generalisation, that is given an unknown example predict its class label with high accuracy. This schema was repeated for all 5 datasets, and for the SVM and *k*-NN classifiers, trained on (i) LOPIT alone, and (ii) GO CC alone, for comparison with the TL algorithms.

For simplicity, throughout this manuscript we refer to the mouse pluripotent embryonic stem cell dataset as the ‘mouse dataset’, the human embryonic kidney fibroblast dataset as the ‘human dataset’, the *Drosophila* embryos dataset as the ‘fly dataset’, the *Arabidopsis thaliana* callus dataset as the ‘callus dataset’ and finally the second *Arabidopsis thaliana* roots dataset, as the ‘roots dataset’.

## The *k*-NN transfer learning classifier

The median macro-F1 scores for the mouse, human, callus, roots and fly datasets were 0.879, 0.853, 0.863, 0.979, 0.965, respectively, for the combined *k*-NN transfer learning approach. A two sample t-test showed that over 100 test partitions, the mean estimated generalisation performance for the *k*-NN transfer learning approach was significantly higher than on profiles trained solely from only primary or auxiliary alone for the mouse (*p* = 2*e*^−21^ for primary alone and *p* = 7*e*^−78^ for auxiliary alone), human (*p* = 1*e*^−7^ for primary alone and *p* = 8*e*^−32^ for auxiliary alone), plant roots (*p* = 4*e*^−17^ for primary alone and *p* = 4*e*^−22^ for auxiliary alone), and fly (*p* = 3*e*^−5^ for primary alone, *p* = 1*e*^−112^ for auxiliary alone) data (Fig. 1). We found that the plant callus dataset did not significantly benefit (nor detrimentally affected) by the incorporation of auxiliary data. This was unsurprising as this dataset is extremely well-resolved in LOPIT (S1 File, Fig. 1, top right) and the median macro F1-score over 100 rounds of training and testing with a baseline *k*-NN classifier resulted in a median macro F1-score of 0.985 (the combined approach yielded a macro F1-score of 0.973).

**Figure 1:**
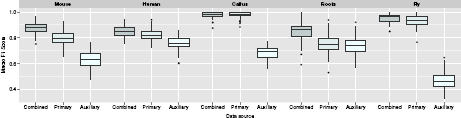
**Boxplots, displaying the estimated generalisation performance over 100 test partitions**. Results for the *k*-NN transfer learning algorithm applied with (i) optimised class-specific weights (combined), (ii) only primary data and (iii) only auxiliary data, for each dataset.

The *k*-NN transfer learning classifier uses optimised class weights to control the proportion of primary to auxiliary neighbours to use in classification. One advantage of this approach is the ability for the user to set class weights manually, allowing complete control over the amount of auxiliary data to incorporate. As previously described, the class weights can be set through prior optimisation on the labelled training data. Fig. 2 shows the detailed results for the mouse dataset and the distribution of the 100 best weights selected over 100 rounds of optimisation are shown on the top left. We found the distribution of weights for each dataset reflected closely the sub-cellular resolution in each experiment. For example, for the experiment on the mouse dataset the distribution of best weights identified for the endoplasmic reticulum (ER), mitochondria and chromatin niches are heavily skewed towards 1 indicating that the proportion of neighbours to use in classification should be predominantly primary. Note, as described in the methods if the class weight is assigned to 1, then strictly only neighbours in primary data are used in classification and similarly, if the class weight is 0 then all weight is given to the auxiliary data. If the weight falls between these two limits the neighbours in both the primary and auxiliary data sources is considered. From examining the principal components analysis (PCA) plot (Fig. 2, top right) we indeed found that these organelles are well separated in the LOPIT experiment. Conversely, we found that the 40S ribosome overlaps somewhat with the nucleus (non-chromatin) cluster (Fig. 2, top right) which is reflected in the best choice of class weights for these two niches; they were both assigned best weights of 1/3 and their weight distributions are skewed towards 0 indicating that more auxiliary data should be used to classify these sub-cellular classes. If we further examine the class-F1 scores for these two sub-cellular niches (Fig. 2, bottom) we indeed find that including the auxiliary data in classification yields a significant improvement in generalisation accuracy (*p* = 1*e*^−16^ for 40S ribosome (red) and *p* = 1*e*^−10^ for the nucleus (non-chromatin) (pink)). We also found this to be the case for the proteasome, which is overlapping with the cytosol. We found LOPIT alone did not distinguish between these two sub-cellular niches in this particular experiment, however, the addition of auxiliary data from the Gene Ontology resulted in a significant increase in classifier prediction (*p* = 2*e*^−16^) as shown by the class-specific box plot in Fig. 2, bottom (black).

**Figure 2:**
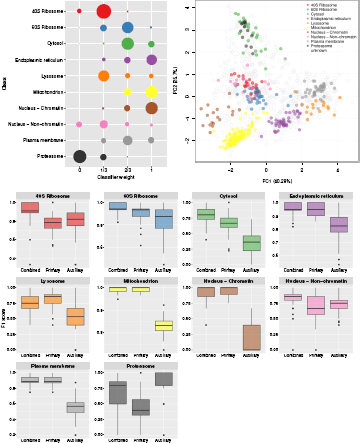
**Visualisation of *k*-NN TL results**. Top left: Bubble plot, displaying the distribution of the optimised class weights over the 100 test partitions for the transfer learning algorithm applied to the mouse dataset. Top right: Principal components analysis plot (first and second components, of the possible eight) of the mouse dataset, showing the clustering of proteins according to their density gradient distributions. Bottom: Sub-cellular class-specific box plots, displaying the estimated generalisation performance over 100 test partitions for the transfer learning algorithm applied with (i) optimised class-specific weights (combined), (ii) only primary data and (iii) only auxiliary data, for each sub-cellular class.

Many experiments are specifically targeted towards resolving a particular organelle of interest (e.g. the TGN in the roots dataset) which requires careful optimisation of the LOPIT gradient. In such a setup sub-cellular niches other than the one of interest may not be well-resolved which may simply be due to the fact that the gradient was not optimised for maximal separation of all sub-cellular niches, but only one or a few particular organelles. Such experiments in particular may benefit from the incorporation of auxiliary data. We found that for the roots dataset all sub-cellular classes, except the TGN sub-compartment, benefitted from including auxiliary data (S1 File Fig. 3, bottom), highlighting the advantage of using more than one source of information for sub-cellular protein classification. The best weight for the TGN was found to be 1 (S1 File Fig. 3, top left), as expected and indicating high resolution in LOPIT for this class. In this framework we are able to resolve different niches in the data according to different data sources, further highlighted in the class-specific boxplots in S1 File Figs 1 to 4.

## The SVM transfer learning classifier

Adapting Wu and Dietterich’s classic application of transfer learning [6] we have implemented an SVM transfer learning classifier that allows the incorporation of a second auxiliary data source to improve upon the classification of experimental and condition-specific sub-cellular localisation predictions. The method employs the use of two separate kernels, one for each data source. As previously described, to assess the generalisation accuracy of our classifier we employed the classic machine learning schema of partitioning our labelled data into training and testing sets, and used the testing sets to assess the strength of our classifiers. This was repeated on 100 independent partitions for (i) the SVM TL method, (ii) a standard SVM trained on LOPIT alone, and (iii) a standard SVM trained on GO CC alone.

For the SVM TL experiments the resultant median macro-F1 scores for the mouse, human, callus, roots and fly datasets were 0.902, 0.868, 0.956, 0.875, 0.961, respectively, over the 100 partitions. As per the *k*-NN TL, we found the macro-F1 scores for the SVM TL (S2 File Fig. 1) were significantly higher than on profiles trained solely from only primary or auxiliary alone; mouse (*p* = 5*e*^−56^ for primary alone and *p* = 6*e*^−37^ for auxiliary alone), human (*p* = 7*e*^−3^ for primary alone and *p* = 1*e*^−21^ for auxiliary alone), callus (*p* = 4*e*^−3^ for primary alone and *p* = 1*e*^−92^ for auxiliary alone), roots (*p* = 2*e*^−45^ for primary alone and *p* = 7*e*^−25^ for auxiliary alone), and fly (*p* = 3*e*^−3^ for primary alone and *p* = 4*e*^−105^ for auxiliary alone) data. This was also evident on the organellar level as seen in the supporting figures in the S2 File.

## Other auxiliary data sources

One of the advantages of the transfer learning framework is the flexibility to use different types of information for both the primary and auxiliary data source. We demonstrate the flexibility of this framework by testing other complementary sources of information as an auxiliary data source.

**The Human Protein Atlas**. The sub-cellular Human Protein Atlas [57] provides protein expression patterns on a sub-cellular level using immunofluorescent staining of human U-2 OS cells. As described in the materials and methods the hpar Bioconductor package [60] was used to query the sub-cellular Human Protein Atlas [57] (version 13, released on 11/06/2014). This auxiliary data, to be integrated with our human LOPIT experiment, was encoded as a binary matrix describing the localisation of 670 proteins in 18 sub-cellular localisations. Information for 192 of the 381 labelled marker proteins were available. These 192 proteins covered 8 of the 10 known localisations in the human LOPIT experiment and were used to estimate the classifier generalisation accuracy of (i) the transfer learning approach with both data sources, (ii) the LOPIT data alone and (iii) the HPA data alone, as described previously. As detailed in the supplementary information (S3 File Fig. 1), we observed a statistically significant improvement in our overall classification accuracy as well as several positive organelle-specific results.

**YLoc sequence and annotation features**. Sequence and annotation features, that were used as input from the computational classifier YLoc [58, 59] (see materials and methods, Table 1) were selected as an auxiliary data source to complement the LOPIT mouse stem cell dataset. 34 sequence and annotation features were selected using a correlation feature selection, as described in the materials and methods. Using the LOPIT mouse dataset as our primary data, and the 34 YLoc features as our auxiliary we employed the standard protocol for testing classifier performance (i) using the *k*-NN transfer learning with both data sources, (ii) the primary data alone and (iii) the auxiliary data alone. Although we did not observe a statistically significant improvement using the auxiliary data in the transfer learning framework, we did not see any statistically significant disadvantage in combining information (S3 File Fig. 2). Thus we found that incorporating data from auxiliary sources in this framework does not dilute any strong signals in the original experiment, demonstrating the flexibility of the classifier.

**Protein-protein interaction data**. Protein-protein interaction data was retrieved from the STRING database [54] (version 10) in the human data set. An interaction contingency matrix was constructed using the STRING combined scores (see methods). Interaction scores for 1109 possible interaction partners were available for 99 of the 381 markers. As described for the other sources above, using this protein-protein interaction information as an auxiliary data source we employed the standard protocol for testing classifier performance (i) using the *k*-NN transfer learning with both data sources, (ii) the primary data alone and (iii) the auxiliary data alone. As per the YLoc data we did not observe a statistically significant increase in combining auxiliary information with our primary data using transfer learning, however, we did not see any statistically significant disadvantage (S3 File Fig. 3).

## Biological application

We applied the two transfer learning classifiers to a real-life scenario, using the E14TG2a mouse stem cell dataset as our use-case to (i) demonstrate algorithm application, and (ii) highlight the applicability of the method for predicting protein localisation in MS-based spatial proteomics data over other single-source classifiers.

**Sub-cellular protein localisation prediction in mouse pluripotent embryonic stem cells**. The E14TG2a mouse stem cell LOPIT dataset contained 387 labelled and 722 un-labelled protein protein profiles distributed among 10 sub-cellular niches (Table 1 of the S6 File). Following extraction of the GO CC auxiliary data matrix for all proteins in the dataset the following four classifiers were applied (1) a *k*-NN (with LOPIT data only), (2) the *k*-NN TL (with LOPIT and GO CC data), (3) an SVM (with LOPIT data only) and (4) the SVM TL (with LOPIT and GO CC data) and the parameters for each optimised (see methods) for the prediction of the sub-cellular localisation of the unlabelled proteins in the dataset.

In supervised machine learning the instances which one wishes to classify can only be associated to the classes that were used in training. Thus, it is common when applying a supervised classification algorithm, wherein the whole class diversity is not present in the training data, to set a specific score cutoff on which to define new assignments, below which classifications are set to unknown/unassigned. The `pRoloc` tutorial, which is found in the set of accompanying vignettes in the `pRoloc` package [56], describes this procedure and how to implement this in practice. Deciding on a threshold is not trivial as classifier scores are heavily dependent upon the classifier used and different sub-cellular niches can exhibit different score distributions.

To validate our results and calculate classification thresholds based on a 5% false discovery rate (FDR) for each of the four classifiers (i.e. *k*-NN, *k*-NN TL, SVM, SVM TL) we compared the predicted localisations with the localisation of the same proteins found in the highest resolution spatial map of mouse pluripotent embryonic stem cells to date [61]. From examining the overlap between our new classifications and the localisations in the high resolution mouse map we found 183 of our 722 unlabelled proteins matched a high confidence localisation in the new dataset. Of the remaining, 347 of our proteins were labelled as unknown in the mouse map (i.e. were assigned a low confidence localisation in the experiment), and 192 proteins did not appear in the map. We used the localisation of these 183 high confidence proteins as our gold standard on which to validate our findings and set a FDR for our predictions.

**Increasing classifier discrimination power**. S4 File Fig. 1 shows the score distributions for correct and incorrect assignments of the unassigned proteins in the dataset (as validated through the hyperLOPIT mouse map [61]) and the distribution of the scores per classifier. Note, the scores are not a reflection of the classification power and the score distributions between the four different methods are not comparable to one another as they are calculated using different techniques. For both of the single-source *k*-NN and SVM classifiers there is a large overlap in the distribution of scores for correct and incorrect assignments (S4 File Fig. 1). It is desirable to have a distribution of scores that enables one to choose a cutoff that minimises the FDR. What is evident from examining the score distributions of incorrect and correct assignments is that by using transfer learning we have increased the discrimination power of the classifier and thus lowered our FDR.

This is further highlighted by receiver-operator characteristics (ROC) analysis (Fig. 3) in which the performance of the 4 different classifiers is displayed for different scoring thresholds. When given a specific score cutoff, the ROC curve plots the true-positive rate (TPR) versus the false-positive rate (FPR) for each classifier. We calculated the area under the ROC curve (AUC) for each classifier and found the AUC for the *k*-NN, SVM, *k*-NN TL and SVM TL was 0.693, 0.705, 0.746 and 0.822, for each classifier respectively.

Using our knowledge of the correct/incorrect outcomes of these 183 previously unlabelled proteins we calculated an appropriate threshold at which to classify all unlabelled proteins. Using a FDR of 5% we found assignment thresholds for the SVM (0.85), SVM TL (0.785) and *k*-NN TL (0.805) to classify the remaining unlabelled proteins. A FDR of 5% was not possible with the *k*-NN classifier, and the lowest achievable FDR was 15%, which occurred using the strictest threshold of 1 i.e. only when all 5 nearest neighbours agreed. Comparing the classifications made from the single-source classifiers to those made with the transfer learning methods, we found in both cases we get many more assignments using the combined transfer learning approaches compared to the single-source methods using a fixed FDR of 5%, as discussed below.

Fig. 4 shows the SVM and SVM TL scores assigned to each of the 183 validated proteins.

**Figure 3:**
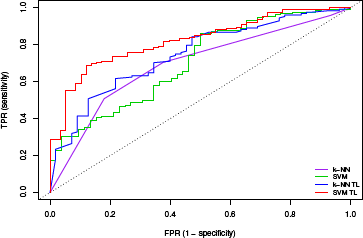
**Receiver-operator characteristics (ROC) analysis**. The performance of the 4 different algorithms for varying scoring thresholds. For a specific score cutoff, the ROC curve shows the true-positive rate (TPR) versus the false-positive rate (FPR) for each classifier. We calculated the area under the ROC curve (AUC) for each curve; the AUC for the *k*-NN, SVM, *k*-NN TL and SVM TL classifiers were 0.693, 0.705, 0.746 and 0.822, for each classifier respectively.

The sub-cellular class is highlighted by solid colours and an un-filled point on the plot represents the case where the two classifiers disagreed on the sub-cellular localisation. We found that the SVM TL classifier gave 70% more high-confidence classifications with the same 5% FDR threshold than the single-source SVM trained on primary data alone. All proteins that were assigned to a sub-cellular niche with a high confidence score in both the SVM and SVM TL (Fig. 4, top right grid) were assigned to the same class. We also found that many proteins outside of the high confidence threshold were assigned the same sub-cellular class using both methods, as indicated by the abundance of solid points on the plot. Of the total 722 previously unlabelled proteins we assigned high confidence localisations for 204 proteins using the SVM TL, and 176 proteins using the *k*-NN TL method, based on a FDR of 5% (Tables 1 and 2 of the S4 File).

**New findings**. By way of biological validation we investigated the additional protein assignments that were found using the SVM TL method (Fig. 4, bottom right grid) as novel assignments to one of these classes, the plasma membrane, by searching through the literature for supporting empirical evidence. For example, using the SVM TL method we found four new proteins (GTR3 MOUSE, SNTB2 MOUSE, PAR6B MOUSE and ADA17 MOUSE) assigned only to the plasma membrane with the SVM TL method (Fig. 5) that were also assigned to the plasma membrane in the recent high resolution mouse map [61] (S4 File Fig. 2). Dehydroascorbic acid transporter (GTR3 MOUSE) is a multi-pass membrane protein which has been previously shown to be a plasma membrane protein in studies isolating the cell surface glycoprotein in Jurkat cells [62]. Beta-2 syntrophin or syntrophin 3 (SNTB2 MOUSE) is a phosphoprotein with PDZ domain through which it interacts with ion channels and receptors. There are confounding reports of the sub-cellular location of this peripheral protein. It associates with dystrophins and has no signal sequence. It is found mostly in muscle fibres and brain [63], but to date, its role has not been studied in mouse embryonic stem cells. Given its association with ion channels and receptors, it is perfectly feasible that the steady location of this protein in stem cells is the plasma membrane. Partitioning defective 6 homolog beta (PAR6B MOUSE) is a peripheral membrane protein thought to be in complex with E-cadherin, aPKC, and Par3 at the plasma membrane [64], where it functions to guide GTP-bound Rho small GTPases to atypical protein kinase C proteins [65]. Disintegrin and metalloproteinase domain-containing protein 17 (ADA17 MOUSE) is a single pass plasma membrane protein which functions to cleave the intracellular domain of various plasma membrane proteins including notch and TNF-alpha [66]. It is therefore involved in the upstream events in several signalling pathways. It has a 17 amino acid N-terminal signal sequence suggestive of its function as a membrane protein. The full list of localisation predictions for all proteins in the mouse dataset can be found in the R data package `pRolocdata`.

**Figure 4:**
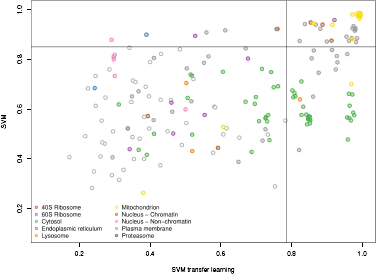
Scatterplot displaying the scores for the SVM and SVM TL classifiers for the 183 proteins validated by the hyperLOPIT mouse map [61]. Each point represents one protein and its associated classifier scores. Filled circles highlight proteins that were assigned the same sub-cellular class with each classifier, empty circles represent the instance when the two classifiers gave different results. The solid lines show the classification boundaries for the two classifiers at a 5% FDR, above which proteins are classified to the highlighted class, below these boundaries proteins are deemed low confidence and thus left unassigned.

**Figure 5:**
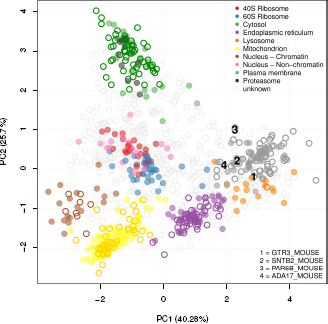
**Principal components analysis plot (PCA) of the mouse stem cell dataset**. Proteins are clustered according to their density gradient distributions. Each point on the PCA plot represents one protein. Filled circles are the original protein markers used in classification, hollow circles show new locations as assigned by the SVM TL classifier. The 4 proteins GTR3 MOUSE, SNTB2 MOUSE, PAR6B MOUSE and ADA17 MOUSE that were found in the SVM TL method and not in an SVM classification with LOPIT only are highlighted.

## A Comparison of Transfer Leaning Algorithms

We compared the macro- and class-F1 scores from all experiments on all 5 datasets used to assess the classifier performance of the *k*-NN TL and SVM TL methods. We found that no single method systematically outperformed the other, as described in the S5 File of the supporting supplement.

When applying the SVM TL and *k*-NN TL classifiers to the unlabelled proteins (see biological validation) an analysis of the final assignments (as classified based on a FDR of 5%) showed that the predicted protein localisations were in high agreement. Although there were no protein-organelle assignment mismatches between TL methods we did find a few cases where one TL method would assign a protein to one of the sub-cellular classes but the other TL method did not result in any organelle assignment, due to low classification scores (see S4 File Table 3). Overall, we did not find any contradicting sub-cellular class assignments.

We also compared Wu’s original *k*-NN algorithm against our class-specific implementation (see S5 File Figure 6). Wu’s method was better than using primary data alone for all but the callus dataset, but was significantly outperformed by our method for the mouse (p = 4*e*^−4^) and roots dataset (p = 4*e*^−3^).

## Discussion

In this study we have presented a flexible transfer learning framework for the integration of heterogeneous data sources for robust supervised machine learning classification. We have demonstrated the biological usage of the framework by applying these methods to the task of protein localisation prediction from MS-based experiments. We further show the flexibility of the framework by applying these methods to the five different spatial proteomics datasets, from four different species, in conjunction with three different auxiliary data sources to classify proteins to multiple sub-cellular compartments. We find the two different classifiers— the *k*-NN TL and SVM TL—perform equally well and importantly both of these methods outperform a single classifier trained on each single data source alone. We further applied the algorithm to a real-life use case, to classify a set of previously unknown proteins in a spatial proteomics experiment on mouse embryonic stem cells, which was validated using the most high resolution map of the mouse E14TG2a stem cell proteome produced to date [61]. We find integrating data from a second data source directly into classifier training and classifier creation results in the assignment of proteins to organelles with high generalisation accuracy. Finally, we find that using freely available data from repositories we can improve upon the classification of experimental and condition-specific protein-organelle predictions in an organelle-specific manner.

To our knowledge, no other method has been developed to date that allows the incorporation of an auxiliary data source for the primary task of predicting sub-cellular localisation in spatial proteomics experiments. In this study we have developed methods that not only allow the inclusion of an auxiliary data source in localisation prediction, but we have created a flexible framework allowing the use of many different types of auxiliary information, and furthermore allowing the user complete control over the weighting between data sources and between specific classes. This is extremely important for the analysis of biological data in general, and spatial proteomics data in particular, as many experiments are targeted towards resolving specific biologically relevant aspects (sub-cellular niches in spatial proteomics) and thus users may wish to control the impact of auxiliary information for aspects that have been specially targeted for analysis by the primary experimental method. In this context the setting of weights manually in the *k*-NN transfer learning classifier allows users complete power to explicitly choose whether to call upon an auxiliary data source or simply use data from their own experiment, on an organelle-by-organelle basis.

The effectiveness of using databases as an auxiliary data source will depend greatly on abundance and quality of annotation available for the species under investigation. For example, human is a well-studied species and there is a large amount of information available in the Gene Ontology and Human Protein Atlas. Furthermore, some organelles are easier to enrich for and thus there exists much more information available to utilise as an auxiliary source on a organelle by organelle basis. The transfer learning methods we present here allow the inclusion of any type of auxiliary data, provided of course there is information available for the proteins under investigation.

The integration of auxiliary data sources is a double-edged sword. On the one hand, it can shed light on (i) the primary classification task by reinforcing weak patterns or (ii) complement the signal in the primary data. On the other hand however it is easy to dilute valuable signals in an expensive experiment by shadowing the uniqueness, and hence biological relevance of the experimental primary data when integration is not performed with care, a phenomenon coined negative transfer (see S5 File Figure 7). Thus one needs to be cautious with data integration in general and not overlook the biological relevance of the primary data. Here, we provide a solution to this issue by using transfer learning: the *k*-NN transfer learning classifier uses optimised class-specific weights so as not to penalise any strong signals in the primary, if no signal is found in the auxiliary; similarly, the SVM transfer learning method uses optimised data-specific gamma parameters for each data-specific kernel.

The transfer learning framework forms part of the open-source open-development Bio-conductor `pRoloc` suite of computational methods available for organelle proteomics data analysis. Moreover, as the pipeline utilises the formal Bioconductor classes, different data types, for example from gene expression technologies among others, can be easily used in this framework. The integration of different data sources is one of the major challenges in the data intensive world of computational biology, and here we offer a flexible and powerful solution to unify data obtained from different but complimentary techniques.

## Materials and Methods

### Primary data

Five datasets, from studies on *Arabidopsis thaliana* [7, 15], *Drosophila* embryos [17], human embryonic kidney fibroblast cells [20], and mouse pluripotent embryonic stem cells (E14TG2a) (unpublished) were collected using the standard LOPIT approach as described by Sadowski et al. [12]. In the LOPIT protocol, organelles and large protein complexes are separated by iodixanol density gradient ultracentrifugation. Proteins from a set of enriched sub-cellular fractions are then digested and labelled separately with iTRAQ or TMT reagents, pooled, and the relative abundance of the peptides in the different fractions is measured by tandem MS. The number of measurements obtained per gradient occupancy profile (which comprises of a set of isotope abundance measurements) is thus dependent on the reagents and LOPIT methodology used.

The first *Arabidopsis thaliana* dataset [7] on callus cultures employed dual use of four isotopes across eight fractions and thus yielded 8 values per protein profile. The aim of this experiment was to resolve Golgi membrane proteins from other organelles. Gradient-based separation was used to facilitate this, including separating and discarding as much nuclear material as possible during a pre-centrifugation step, and carbonate washing of membrane fractions to remove peripherally associated proteins, thereby maximising the likelihood of assaying less abundant integral membrane proteins from organelles involved in the secretory pathway.

The second *Arabidopsis thaliana* dataset on whole roots is one of the replicates published by Groen et al. [15], which was set up to identify new markers of the trans-Golgi network (TGN). The TGN is an important protein trafficking hub where proteins from the Golgi are transported to and from the plasma membrane and the vacuole. The dynamics of this organelle are therefore complex which makes it a challenge to identify true residents of this organelle. For each replicate, sucrose gradient fractions were subjected to a carbonate wash to enrich for membrane proteins and four fractions were iTRAQ labelled. Following MS the resultant iTRAQ reporter ion intensities for the four fractions were normalised to six ratios and then each protein’s abundance was further normalised across its six ratios by sum. In Groen’s original experiment the iTRAQ quantitation information for common proteins between the three different gradients were concatenated to increase the resolution of the TGN [23].

The aim of the *Drosophila experiment* [17] was to apply LOPIT to an organism with heterogeneous cell types. Tan et al. [17] were particularly interested in capturing the plasma membrane proteome (personal communication). There was a pre-centrifugation step to deplete nuclei, but no carbonate washing, thus peripheral and luminal proteins were not removed. In this experiment four isotopes across four distinct fractions were implemented and thus yielded four measurements (features) per protein profile.

The human dataset [67, 20] was a proof-of-concept for the use of LOPIT with an adherent mammalian cell culture. Human embryonic kidney fibroblast cells (HEK293T) were used and LOPIT was employed with 8-plex iTRAQ reagents, thus returning eight values per protein profile within a single labelling experiment. As in the LOPIT experiments in *Arabidopsis* and *Drosophila*, the aim was to resolve the multiple sub-cellular niches of post-nuclear membranes, and also the soluble cytosolic protein pool. Nuclei were discarded at an early stage in the fractionation scheme as previously described, and membranes were not carbonate washed in order to retain peripheral membrane and lumenal proteins for analysis.

The E14TG2a embryonic mouse dataset (unpublished) also employed iTRAQ 8-plex labelling, with the aim of cataloguing protein localisation in pluripotent stem cells cultured under conditions favouring self-renewal. In order to achieve maximal coverage of sub-cellular compartments, fractions enriched in nuclei and cytosol were included in the iTRAQ labelling scheme, along with other organelles and large protein complexes as for the previously described datasets. No carbonate wash was performed.

For validation of the predicted localisations made using the transfer learning classifiers on the E14TG2a dataset above, a new high resolution mouse map was used as a gold standard [61]. This high resolution map was generated using hyperplexed LOPIT (hyperLOPIT), a novel technique for robust classification of protein localisation across the whole cell. The method uses an elaborate sub-cellular fractionation scheme, enabled by the use of Tandem Mass Tag (TMT) 10-plex and application of a recently introduced MS data acquisition technique termed synchronous precursor selection MS3 (SPS)-MS^3^ [68], for high accuracy and precision of TMT quantification. The study used state-of-the-art data analysis techniques [67, 56] combined with stringent manual curation of the data to provide a robust map of the mouse pluripotent embryonic stem cell proteome. The authors also provide a web interface to the data for exploration by the community through a dedicated online R shiny [69] application (https://lgatto.shinyapps.io/christoforou2015).

All datasets are freely distributed as part of the Bioconductor [55] `pRolocdata` data package [56].

## Auxiliary data

**The Gene Ontology**. The Gene Ontology (GO) project provides controlled structured vocabulary for the description of biological processes, cellular compartments and molecular functions of gene and gene products across species [4]. For each protein seen in every LOPIT experiment the protein’s associated Gene Ontology (GO) cellular component (CC) namespace terms were retrieved using the `pRoloc` package [56]. Given all possible GO CC terms associated to the proteins in the experiment we constructed a binary matrix representing the presence/absence of a given term for each protein, for each experiment.

**Human Protein Atlas**. The Human Protein Atlas (HPA) [57] (version 13, released on 11/06/2014) was used as an auxiliary source of information to complement the human LOPIT dataset. The sub-cellular HPA provides protein expression patterns on a sub-cellular level using immunofluorescent staining of human U-2 OS cells. We used the hpar Biocon-ductor package [60] to query the atlas. The data was encoded as a binary matrix describing the localisation of 670 proteins in 18 sub-cellular localisations. In the HPA the reliability of annotated protein expression data is given a status of supportive or uncertain, dependent on similarity to immunostaining patterns and consistency with available experimental gene/protein characterisation data in the UniProtKB database. Here, we only localisations that have been supportively identified.

**YLoc Classifier Features**. YLoc [58, 59] is an interpretable web server developed by Briesemeister and co-workers for the prediction of protein sub-cellular localisation. The YLoc classifier uses numerous features derived from sequence and annotation. A summary of the features included in the YLoc classifier is shown in Table 1. These features provide a source of complementary auxiliary data for the high quality MS based datasets described above. To use these features as an auxiliary source of information, a large-scale correlation-based feature selection (CFS) approach [70], as described in [58, 59], was used with the markers from the mouse dataset to find the set of the most important features.

**Protein-protein interaction data**. The STRING (Search Tool for the Retrieval of Interacting Genes/Proteins) database [54] contains known and predicted protein interactions and quantitatively integrates interaction data from direct (physical) and indirect (functional) associations for a large number of organisms, including human. We have queried the STRING database (version 10) with protein accessions and retrieved the interaction partners of proteins in the human LOPIT data. For each of these proteins, an interaction was recorded and scored using the STRING combined interaction score which was then used to construct an interaction contingency matrix to use as an auxiliary data source. For the 1371 proteins in our human dataset, 520 proteins (99 markers) displayed interactions, which were used in classifier testing.

The creation of the auxiliary datasets are documented and demonstrated using executable code in the *pRoloc-transfer-learning* vignette.

**Table 1:**
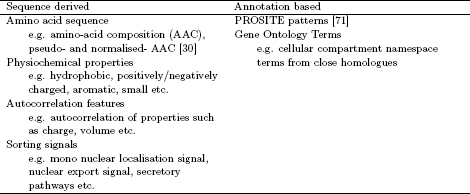
A summary of the types of features considered in training and building Briesemeister et al’s YLoc classifier.

The definition of primary and auxiliary is not defined algorithmically by the quality or the size of the data, but rather by the data and question at hand. Here, LOPIT was considered the primary data because it represented the experiment of interest that was to be complemented by the auxiliary data. In fact, from an algorithmic point of view, primary and auxiliary are reciprocal.

## Markers

Spatial proteomics relies extensively on reliable sub-cellular protein markers to infer proteome wide localisation. Markers are proteins that are defined as reliable residents and can be used as reference points to identify new members of that sub-cellular niche. Here, marker proteins are selected by domain experts through careful mining of the literature. Markers for each LOPIT experiment were specific to the system under study and conditions of interest and are distributed as part of the Bioconductor [55] `pRoloc` package [56].

## Notation

The primary MS-based experimental datasets *P* consist of multivariate protein profiles. The auxiliary data *A* is a presence/absence binary matrix of Gene Ontology Cellular Compartment (GO CC) terms. Data are annotated to either (i) a single known organelle (labelled data), or (ii) have unknown localisation (unlabelled data). Thus we split *P* and *A* into labelled (*L*) and unlabelled (*U*) sections such that *P* = (*L^P^*, *U^P^*) and *A* = (*L^A^*, *U^A^*).

The labelled examples for *P* and *A* are represented by *L^P^* = {(**x**_*l*_,_*yl*_)|*l* = 1,…,|*L^P^*|} where **x**_*l*_ ∈ ℝ^*S*^, and *L^A^* = {(**v**_*l*_,*y_l_*)|*l* = 1,…, |*L^A^*|} where **v**_*l*_ ∈ ℝ^*T*^. Thus each *l*^th^ protein is described by vectors of *S* and *T* features (generally, *S* << *T*), for *P* and *A* respectively. Each dataset shares a common set of proteins that is annotated to one of the same *y_l_* ∈ *C* = {1,…, |*C*|} sub-cellular classes, where |*C*| ∈ ℕ is the total number of sub-cellular classes. Unlabelled data, *U^P^* and *U*^A^ are represented by *U^P^* = {**x**_*u*_|*u* = 1,…, |*U^P^*|} where **x**_*u*_ ∈ ℝ^*S*^ and *U^A^* = {**v**_*u*_|*u* = 1,…, |*U^A^* |} where **v**_*u*_ ∈ ℝ^T^, respectively.

The labelled data for the *i*^th^ organelle class, with *N_i_* indicating the number of proteins for the *i*^th^ organelle class, is given for *P* by 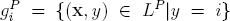 and for *A* by 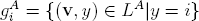. The labelled dataset of all available proteins over the |*C*| different sub-cellular classes is given for *P* by 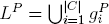 and for *A* by 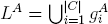.

## Transfer learning using a *k*-nearest neighbours framework

We adapt Wu and Dietterich’s [6] classic application of inductive transfer using experimental quantitative proteomics data as the primary source (*P*) and GO CC terms as the auxiliary source (*A*). We aim to exploit auxiliary data to improve upon the sub-cellular classification of proteins found in MS-based LOPIT experiments in an organelle-specific way, using the baseline *k*-nearest neighbours (*k*-NN) algorithm in a transfer learning framework.

In *k*-NN classification, an unknown example is classified by a majority vote of its labelled neighbours, with the example being assigned to the class most common among its *k* nearest neighbours. Independent of the transfer learning classifier we compute the best *k* for each data source for values *k* ∈ {3, 5, 7, 9,11,13,15} through an initial 100 rounds of 5-fold cross-validation using each set of labelled training data for *P* and then independently for *A* (as implemented in `pRoloc`). We denote by *k_P_* the best *k* for *P*, and by *k_A_* the best *k* for *A*.

Having obtained the best *k* for each data source, the transfer learning algorithm works as follows. For the *u*^th^ protein (**x**_*u*_, **v**_*u*_) we wish to classify in *U*, we start by finding the *k_P_* and *k_A_* labelled nearest neighbours for **x**_*u*_ and **v**_*u*_ in *L^P^* and *L^A^*, respectively. Denote these sets 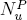 and 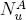. We then define the vectors 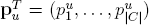 and 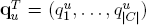 to contain counts for each class in the sets of nearest neighbours; that is,

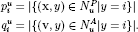

For each protein, let 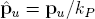 and 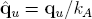 be normalized vectors with elements summing to 1 and representing the distribution of classes among the sets of nearest neighbours for each protein. Finally, let 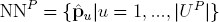 and 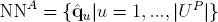.

To include both the primary and auxiliary data in the set of potential neighbours we took a weighted combination of the votes in NN^*P*^ and NN^*A*^ for each sub-cellular class. Class weights are defined by the parameter vector ***θ**^T^* = (*θ*_1_,…,*θ*_|*C*|_) with values 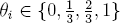 chosen by optimisation through a prior 100 independent rounds of 5-fold cross-validation on a separate training partition of the labelled data. For the *u*^th^ unknown protein (**x**_*u*_, **v**_*u*_) in *U*, the voting scores for each class *i* ∈ *C* are calculated as

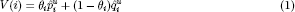

and the protein is assigned to the class *c* ∈ *C* maximizing *V(i*)

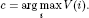

The class weights *θ_i_* in equation 1 control the relative importance of the two types of neighbours for each class *i* ∈ *C*. This differs from Wu and Dietterich’s [6] original approach as they only weight the data sources and not the classes and the data sources. In this paper we select each class weight *θ_i_* from the set 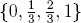 however, the algorithm allows us to use any real-valued *θ_i_* ∈ [0,1]. If *θ_i_* = 1, then all weight is given to the primary data in class *i* and only primary nearest neighbours in class *i* are considered. Similarly, if *θ_i_* = 0, then all weight is given to the auxiliary data in class *i* and only auxiliary nearest neighbours in class *i* are considered. If 0 < *θ_i_* < 1 then a combination of neighbours in the primary and auxiliary data sources is considered.

## Transfer learning using an SVM framework

**Linear programming SVMs** The method is based on the use of the linear programming formulation of the SVM (lpSVM). This formulation promotes classifiers that are sparse, in the sense that where possible only a few parameters obtained through training are non-zero; for a detailed introduction see Mangasarian [72].

We begin by describing the standard lpSVM used for classical two-class classification problems with a single labelled training set. We use the multiple-class version of this approach with the individual primary and auxiliary sets *P* and *A* as a comparison later in the paper; we present the method here assuming that the primary set *P* is being used and can be set up as a binary classification problem; for example, we might wish to predict whether or not a protein should be assigned to a single specified sub-cellular class. For binary classification problems with class labels *y* ∈ {+1,−1}, and given labelled data *L^P^* = {(**x***_l_*,y_*l*_)|*l* = 1,…,*m*} where *m* = |*L^P^*| the classifier takes the form

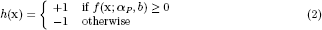

where f is the latent function

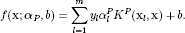

Here, *K^P^* is a kernel (Shawe-Taylor and Cristianini [73]) associated with the primary data and 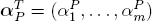 and *b* are parameters determined by training.

For any vector **x**^*T*^ = (*x*_1_,…, *x*_n_) let |.|_1_ denote the 1-norm

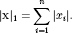

The training algorithm requires that we solve the linear programme

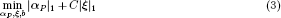

such that for each *i* = 1,…, *m*

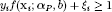

and ***α**_P_,**ξ***, ≥ 0. The parameters ***ξ*** and *C* act in the same way as the corresponding parameters in the standard SVM: ***ξ*** contains the slack variables allowing some examples to be misclassified, and *C* controls the extent to which such misclassifications are penalized during training.

Note that it is possible for the linear programme to have no solution, although we found this to be extremely rare. When this was the case the classifier reverted to predicting the most common class in the labelled data.

**Transfer learning for binary classification**. Once again we adapt the method of Wu and Dietterich [6] to our problem. The original method requires adaptation as it is designed for data having two important differences compared with ours. First, it does *not* require examples in the labelled data sets *L^P^* and *L^A^* to be in correspondence and for corresponding training examples to share the same label. Second it assumes that *P* and *A* share *the same number of features*. While the first of these differences is easily dealt with as our data is a special case that is already covered, the second is more problematic. If we now introduce the labelled auxilliary data *L^A^* = {(**v**_*l*_, *y_l_*)|*l* = 1,…, *m*} a direct application of the approach in [6] requires us to evaluate kernels of the form *K*(**x**,**v**). As *P* and *A* contain data *with different numbers of features* this presents a problem for any SVM-type method, as kernels are usually required to satisfy the Mercer conditions (Mercer [74]), one of which is that they are symmetric, such that *K*(**x**, **x**′) = *K*(**x**′,**x**). While research on the use of asymmetric kernels has appeared—see for example [75]—even if we relax this requirement a kernel is essentially a measure of the similarity of its arguments, and the question arises of how one might sensibly measure the similarity of a protein profile with a presence/absence vector of GO CC terms. This problem does not arise with Wu and Dietterich’s data as the two sets they use have the same dimension and are derived in a way that makes measuring similarity straightforward.

We therefore simplify the original method as follows. We maintain the machinery employed above for the primary data, and introduce a separate kernel *K^A^* and parameter vector ***α**_A_* for the auxilliary data. A vector to be classified now contains both a protein profile **x** and a GO vector **v**. The latent function becomes

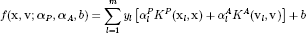

and training requires us to solve the linear program

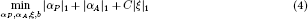

such that for each *i* = 1,…, *m*

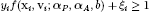

and ***α**_P_,**α**_A_,**ξ***, ≥ **0**.

Note that this differs from the method of *Multiple Kernel Learning (MKL)* (Lanckriet et al. [76], Gönen and Alpaydin [77]) in that in MKL the single kernel *K* is replaced in the usual SVM formulation by a weighted sum of kernels

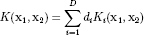

where *d_i_* ≥ 0 and 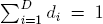. The *d_i_* are then included with ***α*** and *b* in a more involved constrained optimisation problem. Our approach has the advantages that it remains a straightforward linear program and in fact introduces fewer constraints on the form of the latent function *f*.

Throughout our experiments we used for *K^P^* and *K^A^* the Gaussian kernel

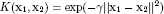

where ||.|| denotes the 2-norm 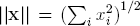. We optimized over the value of *C*, and also separate values *γ_P_* and *γ_A_* for the two kernels as described below, with C in the range {0.125, 0.25,0.5,1, 2, 4, 8,16} and *γ_P_*, *γ_A_* in the range {0.01, 0.1,1,10,100,1000}.

**Multiple classes, class imbalance and probabilistic outputs**. As a baseline comparison in our experiments we used a standard SVM as implemented in the package LIBSVM (Chang and Lin [78]). In extending our transfer learning technique to deal with multiple classes and probabilistic outputs we therefore maintained as close a similarity as possible to the methods used by that library.

SVMs and lpSVMs are in their basic form inherently binary classifiers. In order to address multiple-class problems using non-probabilistic outputs such as the one presented here we use the method of Knerr et al. [79]. We train a binary classifier to separate each pair of classes. In order to classify a new example we then take a vote among these binary classifiers, assigning the example to the class with the most votes.

As we typically have several sub-cellular classes the binary classification problems used in constructing the multiple-class classifier are inherently unbalanced. We adjust for this using the method of Morik et al. [80]. In each binary problem let *n*^+^ denote the number of positive examples and *n*^−^ the number of negative examples. In the linear programme objective functions (equations 3 and 4) we replace the single value for *C* with the adjusted values

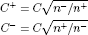

for the positive and negative examples respectively. Let *S*^+^ denote the set of indices of the positive examples and *S*^−^ the set of indices for the negative examples. The term *C*|***ξ***|_1_ in equations 3 and 4 becomes

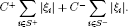

Finally, we prefer to employ probabilistic outputs rather than simply thresholding as in equation 2. Once again we employ the same techniques as LIBSVM. The method for binary classifiers is presented by Platt [81] and Lin et al. [82], and for multiple-class classifiers by Wu et al. [6].

## Assessing classifier generalisation accuracy

In order to evaluate the generalisation accuracy of each transfer learning classifier we employed the following schema in all experiments. A set of LOPIT profiles labelled with known markers, and their counterpart auxiliary GO CC profiles, were separated at random into training (80%) and test (20%) partitions. The split was stratified, such that the relative proportions of each class in each of the two sets matched that of the complete set of data. The test profiles were withheld from classifier training and employed to test the generalisation accuracy of the trained classifiers. On each 80% training partition 5-fold stratified cross-validation was conducted to test all free parameters via a grid search and select the best set of parameters for each classifier. In each experiment, for each dataset, this process of 80/20% stratified splitting, training with 5-fold stratified cross-validation on the 80% and testing on the 20% was repeated 100 times in order to produce 100 sets of macro F1 scores and class-specific F1 scores. The F1 score (He [83]) is a common measure used to assess classifier performance. It is the harmonic mean of precision and recall, where

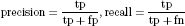

and tp denotes the number of true positives, fp the number of false positives, and fn the number of false negatives. Thus

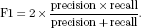

A high macro F1 score indicates that the marker proteins in the test data set are consistently correctly assigned by the algorithm.

To assess whether incorporating an auxiliary data source into classifier training and classifier creation was better than using primary or auxiliary data alone, we conducted three independent experiments for each data source and for each transfer learning method. We used the above schema to assess the generalisation accuracy of using (1) the transfer learning *k*-Nearest Neighbours (*k*-NN) classifier, (2) the primary LOPIT data alone, using a baseline *k*-NN, (3) the auxiliary GO CC data alone, using a baseline *k*-NN. We repeated this for the lpSVM transfer learning classifier and used a standard SVM with an RBF kernel for single data source experiments. Using these experiments we were able to compare using a simple *k*-NN versus the transfer learning *k*-NN, and also the use of a standard SVM versus the combined transfer learning lpSVM approach.

A two-sample two-tailed t-test, assuming unequal variance, was used to assess whether, over the 100 test partitions, the estimated generalisation performance using the optimised class-specific fusion approach was better than using either primary data alone, or auxiliary data alone. A threshold of 0.01 was used in all t-tests to determine significance.

**Optimised parameters for the mouse pluripotent embryonic stem cell data**. To classify the 722 unlabelled proteins in the E14TG2a mouse stem cell dataset we performed 100 rounds of stratified 5-fold cross validation on the training partition as detailed above. The best parameters were found to be *k* = 5 for the *k*-NN classifier and for the *k*-NN TL classifier *k_P_* = 5, *k_A_* = 5 and the best class weights were found to be 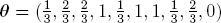 for the 40S ribosome, 60S ribosome, cytosol, endoplasmic reticulum, lysosome, mitochondria, nucleus – chromatin, nucleus – non-chromatin, plasma membrane and proteasome, respectively. For the SVM classifier we found the best parameters to be *C* = 16 and γ = 10. For the SVM TL classifier we found *C* = 16, γ*_P_* = 1 and γ*_A_* = 0.1. Using these parameters with their associated algorithms we classified the 722 unlabelled proteins in the dataset and obtained a classifier score for each protein.

## Supporting Information

### S1 File

**Supporting figures for the *k*-NN transfer learning experiments**. Visualisations of the *k*-NN transfer learning results for human, plant callus, plant roots and fly datasets. Including bubble plots displaying the distribution of the optimised class weights, principal components analysis plots of the LOPIT primary datasets boxplots displaying the estimated generalisation performance of each classifier.

### S2 File

**Supporting figures for the SVM transfer learning experiments**. Boxplots displaying the estimated generalisation performance of the SVM transfer learning algorithm applied to the mouse, human, plant callus, plant roots and fly datasets.

### S3 File

**Supporting figures for other data sources**. Macro-and class-specific results for the *k*-NN transfer learning algorithm used with the auxiliary Human Protein Atlas dataset, a YLoc sequence and annotation features auxiliary dataset and a protein-protein interactions dataset.

### S4 File

**Additional figure and tables for biological application**. Boxplots displaying the distribution of classification scores assigned to the unknown proteins in the mouse dataset for each of the 4 classifiers. Principal components analysis plot displaying the protein classification results from applying the *k*-NN transfer learning algorithm on the unlabelled data in the mouse dataset. Accompanying tables showing the number of sub-cellular assignments of the unlabelled proteins amongst the 10 known sub-cellular classes in the data from applying each transfer learning method. the mouse stem cell dataset highlighting the new localistions found by the *k*-NN transfer learning method.

### S5 File

**A comparison of transfer learning methods**. A short comparison between *k*-NN transfer learning (TL) and SVM TL classifiers, including boxplots displaying the macro-and class-F1 scores for the *k*-NN TL and SVM TL experiments over the 100 test partitions on each dataset. This supplementary file also includes a comparison with Wu’s original *k*-NN transfer learning classifier and negative transfer effects are also described.

### S6 File

**Dataset summary statistics**. Tables displaying the dimensions for each primary and auxiliary dataset, including the total number of proteins identified in each LOPIT dataset and number of known markers of sub-cellular protein location.

## Acknowledgments

LMB was supported by a BBSRC Tools and Resources Development Fund (Award BB/K00137X/1). LG was supported by the European Union 7^*th*^ Framework Program (PRIME-XS project, grant agreement number 262067) and a BBSRC Strategic Longer and Larger Award (Award BB/L002817/1). DW and OK acknowledge funding from the European Union (PRIME-XS, GA 262067) and Deutsche Forschungsgemeinschaft (KO-2313/6-1). The authors would also like to thank Dr. Dureid El-Moghraby from the High Performance Computing Service and Dr. Jenny Barna from Bioinformatics and Computational Biology, University of Cambridge, for their support. This work was performed using the Darwin Supercomputer of the University of Cambridge High Performance Computing Service, provided by Dell Inc. using Strategic Research Infrastructure Funding from the Higher Education Funding Council for England and funding from the Science and Technology Facilities Council.

